# Improving genomic language model reliability under distribution shift

**DOI:** 10.64898/2026.03.19.712858

**Authors:** Gavin Hearne, Mohammad Saleh Refahi, Robi Polikar, Gail L. Rosen

## Abstract

Transformer-based Genomic Language Models (GLMs) have achieved strong performance across diverse genomic prediction tasks. However, their tendency toward overconfident predictions—particularly on noisy or unfamiliar data—limits reliability. In genomics, where unknown species and novel variants are common, developing models robust to distribution shift is crucial for dependable predictions. Here, we analyze the impact of several common and novel uncertainty quantification (UQ) methods in the context of GLMs, evaluating their performance across diverse downstream genomic and metagenomic prediction tasks. Comparing model behavior on both **in-distribution (ID)** and **out-of-distribution (OOD)** data, we show that temperature scaling and epistemic neural networks are capable of improving classification reliability across multiple GLM architectures and domains. The software is available at: https://github.com/EESI/glm-epinet-pyt

## Introduction

In recent years, large language models (LLMs) have risen to become one of the central tools in academia and industry due to their flexibility and ability to learn on unlabeled data [1–3]. In biology, the sequential nature of nucleotide and amino acid sequences makes genomics tasks a suitable application for these tools [4]. Here, genomic sequences are treated as structured, information-rich texts, enabling models to understand genomes in a manner analogous to natural language. These genomic language models (GLMs) are trained on a massive corpus of DNA to predict masked sequences [5–7], building powerful sequence representations capable of diverse downstream predictive tasks, including gene expression prediction, variant effect estimation, and the classification of regulatory elements [8–12]. As a result, GLMs are rapidly becoming a powerful alternative to traditional bioinformatics pipelines, enabling insights into both conserved and species-specific mechanisms of genome function [13, 14].

However, while fine-tuning these models on domain specific tasks has resulted in significant performance gains over both traditional techniques and models trained from scratch, they often fail to accurately assess data that is out of distribution (OOD) [15], resulting in overconfident predictions when applied to new domains [16, 17]. Therefore, it remains an open question: How can we produce more trustworthy genomic AI?

While OOD detection is usually defined as the test data having a different underlying model (or parameters) than that of the training data. Shifts can usually be due to obervational shifts or semantic shifts in the features important to the tasks. Semantic distributions are usually theoretical in the literature – but with biological data, that has inherent evolutionary relations and slow mutational processes, simulating such data becomes more of an issue. Taking already defined datasets and creating our own metagenomic tasks, a contribution of our paper is to define various distributional shifts in genomic data to test reliability of GLMs across tasks.

In comparable deep learning domains, numerous uncertainty quantification (UQ) techniques have been developed to address this by equipping models capable of expressing calibrated confidence and recognizing their own limitations [18–23]. Uncertainty-aware modeling has already shown value in adjacent biological prediction settings, such as microbiome-based prediction, where explicit uncertainty estimates can improve interpretability and downstream decision support [24]. Furthermore, recent work in regulatory genomics has shown that uncertainty-aware training can improve calibration and robustness in genomic deep learning under covariate shift [25]. However, these papers do not compare a variety of uncertainty-aware methods across a variety of tasks. We also create, to the best of our ability, in-distribution (ID), OOD, and varying degrees of “near”-ID and “near”-OOD for the diverse tasks.

In this paper, we compare and contrast both established and recently proposed uncertainty quantification techniques in genomic language modeling applications, to determine how these techniques interact with foundational GLMs across multiple biological domains. Furthermore, we seek to characterize the importance of these techniques within the broader bioinformatics landscape, to understand whether comparable tools are similarly limited in expressing accurate calibration. To do so, we evaluate UQ approaches on 6 downstream classification datasets, encompassing both human cis-regulatory prediction and larger-scale metagenomic gene and taxa classification. We compare the results of these tests across several foundation GLMs, as well as with traditional tools, to establish a baseline for utilizing UQ in the context of genomic language models.

## Methods

### Genomic Language Models

Genomic language models (GLMs), such as DNABERT [6, 7] and Nucleotide Transformer [5], leverage the context-aware capacity of transformer-based networks for genomic sequence modeling. In this study, GLMs are employed as *foundation models*: we adopt publicly available pretrained checkpoints, originally trained on large-scale genomic sequence corpora, and subsequently fine-tune them for specific downstream prediction tasks.

We consider several such foundation models, spanning both transformer-based architectures (Nucleotide Transformer [5], DNABERT-2 [7]) and more novel long-range models (HyenaDNA [9], CARMANIA [26]). All models are fine-tuned for task-specific classification using the Hugging Face `AutoModelForSequenceClassification` API [27].

### Uncertainty Quantification

The need for reliable uncertainty quantification (UQ) has motivated a wide range of methods for estimating neural network confidence. These approaches are often grouped into *deterministic* methods, which produce uncertainty estimates from a single model evaluation [18, 28], and *Bayesian-inspired* methods, which introduce stochasticity or model ensembles to approximate Bayesian model averaging and derive predictive uncertainty [22, 29, 30].

Fig 1 shows one common approximate Bayesian strategy, where multiple stochastic predictive distributions are generated for each input, which can be interpreted as sampling from an approximate posterior over model parameters [19, 31]. Given model parameters *θ*, let 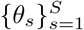 denote *S* stochastic parameter instantiations drawn from *q*(*θ*) ≈ *p*(*θ* | 𝒟) (e.g., independent ensemble members trained on dataset 𝒟). The posterior predictive distribution *p*(*y x*, | 𝒟), given input *x* and label value *y*, is then approximated by the Monte Carlo estimate:

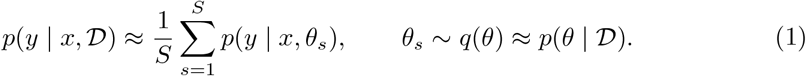

**Fig 1.**
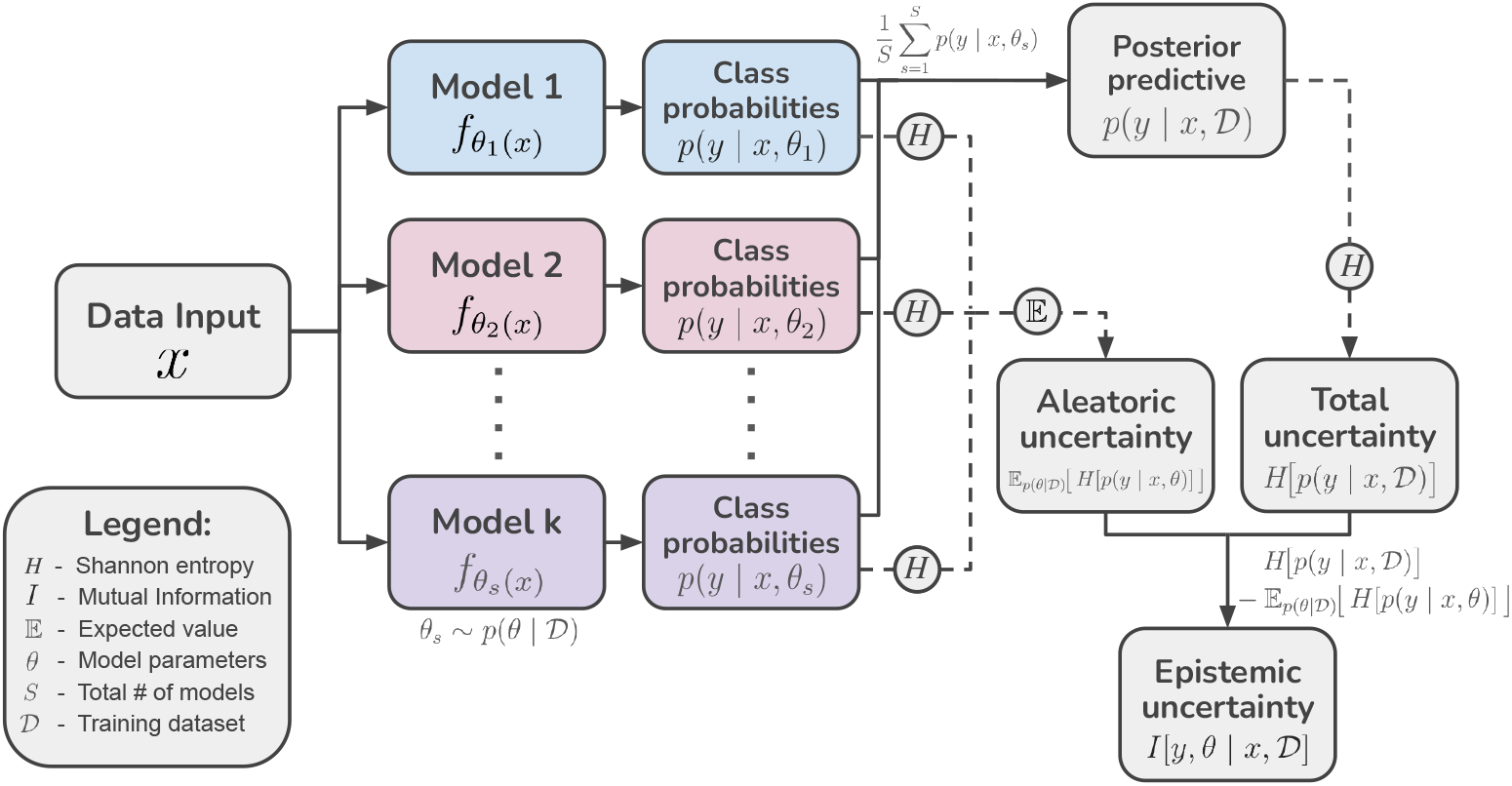
Monte Carlo approximation of the posterior predictive distribution. Approximate Bayesian methods (e.g., deep ensembles, MC dropout, and ENNs/epinets) estimate the posterior predictive distribution by averaging *S* stochastic predictive distributions *p*(*y* | *x, θ*_*s*_) drawn from an approximate posterior *q*(*θ*) ≈ *p*(*θ* | 𝒟).

We then summarize total predictive uncertainty using the predictive entropy *H* of the posterior predictive distribution (where *Y* denotes the random output label) and decompose it into aleatoric and epistemic components via the entropy–mutual information identity [32–34]:

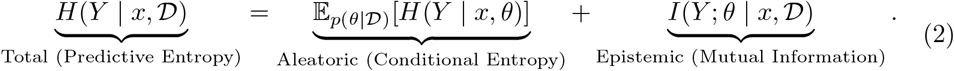

The expected conditional entropy captures *aleatoric* uncertainty(i.e., irreducible uncertainty in the output given the input even under a fixed parameter setting), while the mutual information captures *epistemic* uncertainty, which results from uncertainty over model parameters and is, in principle, reducible with additional data [35–38].

In practice, we estimate these quantities using the same *S* stochastic predictors that define the Monte Carlo posterior predictive above:

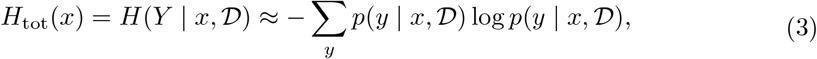

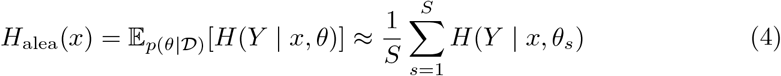

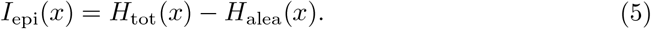

#### Softmax baseline

As a deterministic baseline, we directly use the softmax output of the classification head. From logits **z**, the softmax maps values to probabilities, producing a point estimate of class probabilities **p**. Equation 6 summarizes how predictive uncertainty is approximated by the entropy of this distribution and, where appropriate, by the maximum predicted class probability as a simple confidence score. This baseline does not distinguish between epistemic and aleatoric uncertainty [18, 39].

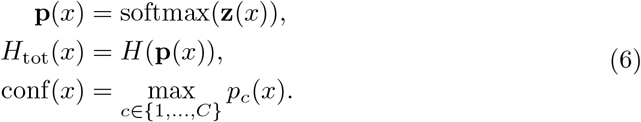

#### Temperature scaling

To better align predicted probabilities with real observed ones, we scale the model output logits by a temperature *T* before applying the softmax [20] as shown in Equation 7. In our experiments, we fit *T* by minimizing the negative log-likelihood on the corresponding held-out validation dataset for each training task. We then fix this value when evaluating on downstream test datasets. Temperature scaling preserves the predicted class label but can reduce overconfidence, improving calibration of probability-based uncertainty scores such as entropy and maximum predicted probability. Like the softmax baseline, temperature scaling remains a deterministic post-hoc method and does not separate epistemic from aleatoric uncertainty.

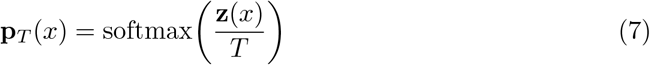

#### Deep ensembles

Deep ensembles approximate uncertainty over model parameters by training *M* independently initialized instances of the same architecture, following Lakshminarayanan et al. [31]. At test time, each ensemble member produces a predictive distribution for a given input, and these distributions are averaged to obtain a marginal predictive distribution (Eq. 8). The spread and disagreement across ensemble members provide a proxy for epistemic uncertainty. While it is generally agreed that deep ensembles are among the strongest baselines for predictive uncertainty [34, 40], the computational cost for both training and inference is prohibitive, preventing practical use for large models [41, 42].

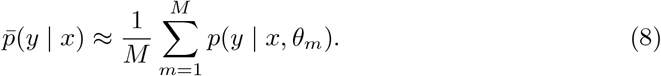

#### Monte Carlo dropout

MC dropout (Fig. 2) implements approximate Bayesian inference by keeping dropout active at test time and performing multiple stochastic forward passes per input, as in Gal et al. [19]. Each pass yields a slightly different predictive distribution due to randomly sampled dropout masks *ξ*; averaging over these predictions produces a Monte Carlo estimate of the marginal predictive distribution as described in Eq. 9. Variation across passes reflects uncertainty in the effective model parameters. Given that a GLM is trained with dropout, implementation is then relatively simple. As opposed to deep ensembles, MC-dropout does not require multiple independent models to be trained. However inference costs are similarly high, and the uncertainty estimations produced by this scheme can often be inaccurate [34, 40, 42, 43].

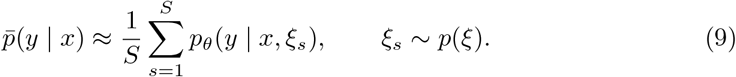

**Fig 2.**
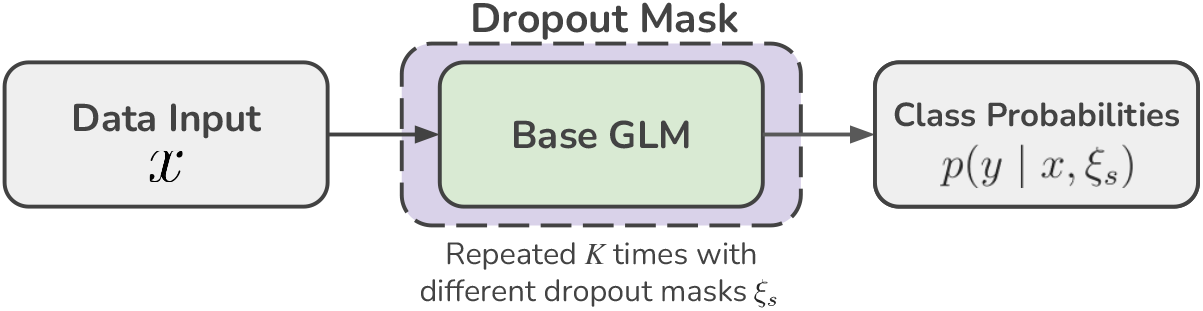
MC-dropout Architecture. Multiple forward passes through the model with different dropout masks, *ξ*_*s*_, produces stochastic outputs, reflecting model uncertainty.

#### Epistemic neural networks (ENNs)

Epistemic neural networks (ENNs) represent epistemic uncertainty by augmenting a predictor with an *epistemic index z* [42]. The index is treated as a latent random variable (drawn from a fixed distribution) that parameterizes a family of predictors *f*_*θ*_(*x, z*), where each value of *z* corresponds to a different plausible hypothesis about the input–output relationship. For a given input *x*, repeated sampling of *z* therefore yields a distribution over predictions, inducing a set of predictive distributions {*p*(*y* | *x, z*)}. Averaging these predictions provides an estimate of the marginal predictive distribution, while variability across index samples provides a principled, ensemble-like summary of epistemic uncertainty without training multiple independent models.

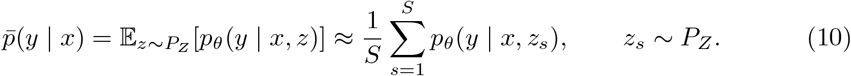

A practical ENN instantiation is the *epinet* [42, 44], which produces index-dependent predictions by combining a shared representation with a lightweight correction head. In our implementation (Fig. 3), the GLM provides a deterministic sequence representation *h*(*x*), and the epinet conditions on *h*(*x*) and a sampled index *z* to generate an *additive* logit correction to the base prediction. The epinet architecture comprises two parts: a *fixed prior* term (a frozen randomized function) that ensures diversity across index samples [45], and a *learnable* component. Through training, the resultant sum of the two networks becomes statistically probable predictions for all values of *z*, while variance in these predictions reflects uncertainty.

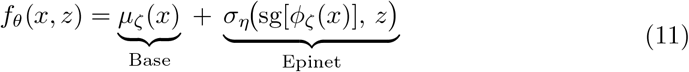

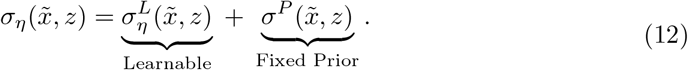

**Fig 3.**
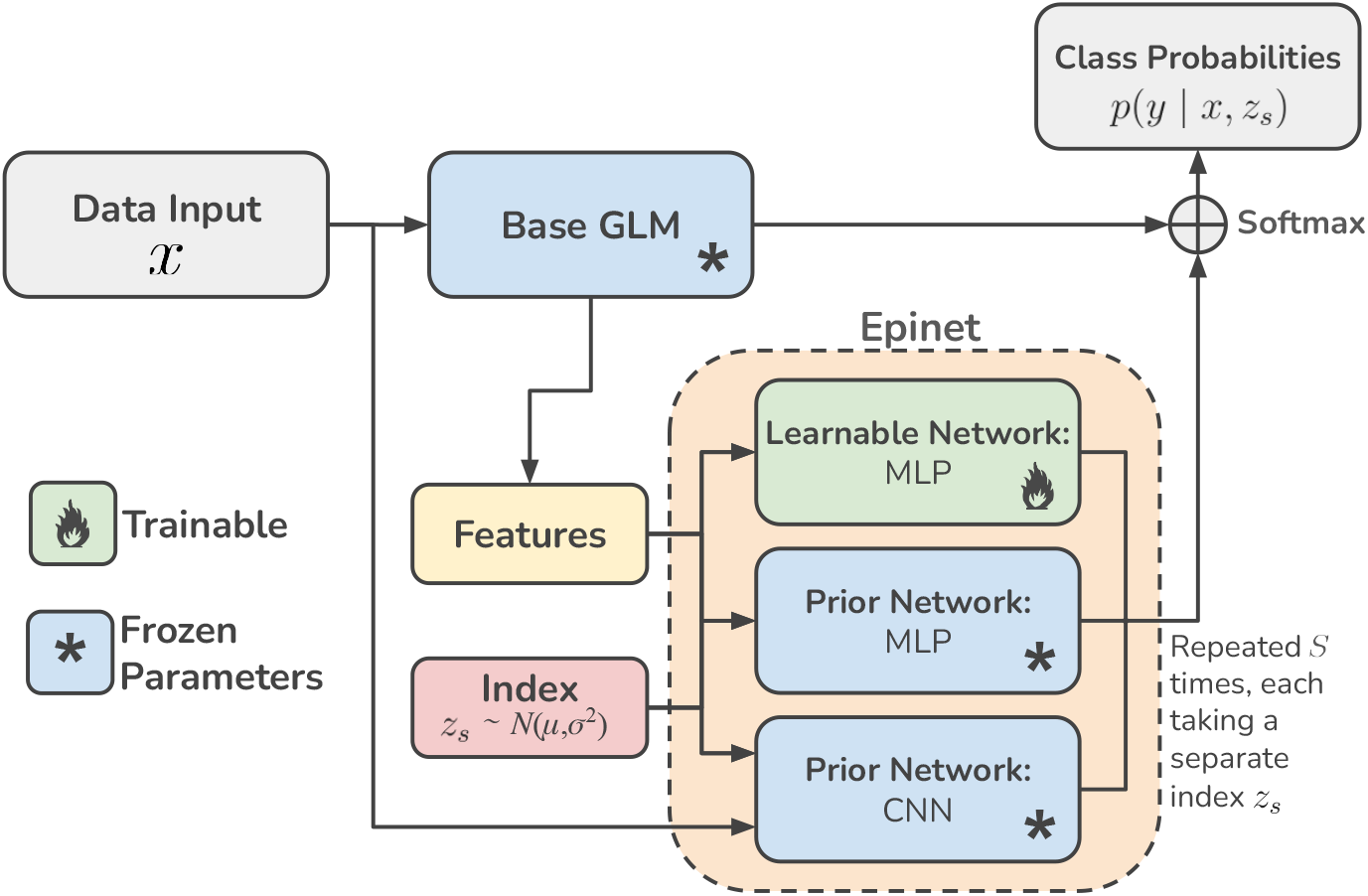
Epinet architecture. The base network produces a standard deterministic prediction along with intermediate features (for a transformer, typically a pooled hidden representation). The epinet takes these features together with a random Gaussian sampled epistemic index *z* and outputs an additive correction to the base prediction, which is applied at the logit level for classification. Repeating inference with different samples of *z* yields a distribution of predictions.

Because the vast majority of GLMs are implemented in PyTorch, we re-implement the core epinet functionality in PyTorch to integrate cleanly with standard GLM training and inference pipelines. We designed the implementation to match the epinet formulation of Osband et al. [42], basing the architecture on the ResNet variant. Here, the epinet prior consists of two components: (i) an MLP prior, matching the learnable network, and (ii) a fixed 1D-convolutional prior ensemble applied directly to the tokenized input. Our code is publicly available at https://github.com/EESI/glm-epinet-pyt. Additional implementation details and verification checks are provided in the Supplementary Material.

### Datasets

We evaluate uncertainty quantification in three biologically distinct regimes: short regulatory element prediction, metagenomic gene classification, and metegenomic taxonomic classification from simulated long reads. These settings were selected because they span different sequence lengths, label spaces, and forms of biological novelty, enabling us to assess calibration and uncertainty under multiple kinds of distribution shift, spanning both functional and taxonomic shift.

Each setting is defined by a fixed training dataset and one or more test datasets chosen to induce increasing degrees of distribution shift, as shown in Table 2. Based on BLAST [46] alignments and biological relatedness, we assign each train–test pair to one of four categories: ID, Near-ID, Near-OOD, and OOD.

#### Regulatory sequence benchmarks

Short regulatory sequence tasks (enhancers, promoters, and splice sites) are derived from the benchmark collection hosted at `InstaDeepAI/nucleotide_transformer_downstream_tasks_revised` [5]. We use their default labeled sequences and train/test splits as the ID condition (e.g., `enhancers_types` trained and evaluated on its own test split). To induce distribution shift while keeping the training set fixed, we evaluate each trained model on held-out test sets from other regulatory tasks in the same benchmark collection. These cross-task evaluations change the task semantics and sequence-function relationship relative to training, producing Near-ID, Near-OOD, or OOD train/test splits.

#### Metagenomic gene classification

The `MsAlEhR/scorpio-gene-taxa` HuggingFace dataset is a gene-and-taxa-classification dataset that includes held-out test sequences intended for evaluating zero-shot generalization performance [11, 47]. For our experiments, we train models to predict gene classes exclusively, using the three provided evaluation conditions to function as different levels of distribution shift: `test`, a held-out split sampled from the same distribution as the training set (ID); `taxa_out` holds out specific taxa present in the original dataset while maintaining the same gene classes (Near-ID); and `gene_out` holds out genes and their associated labels (OOD).

#### Simulated bacterial and non-bacterial reads

For our taxanomic classification setting, we use Pbsim [48] to simulate 6kbp reads from a collection reference genomes sourced from the RefSeq database utilizing Woltka’s pipeline [49, 50], downloaded in July 24, 2023. Models are then trained on bacterial reads for each taxonomic rank (family, order, class, and phylum) and evaluated on three test sets: `novel_genus`, containing reads from previously unseen genera within training families (Near-ID); `novel_family`, containing reads from bacterial families absent from the training set (Near-OOD); and `nonbacterial`, containing reads from non-bacterial taxa (OOD).

### Evaluation protocol

#### Splits and selection

For each task we define 𝒟_train_, 𝒟_val_, and an in-distribution test set 𝒟_test,ID_ using a 90-10 train/validation set. We select checkpoints using 𝒟_val_ according to validation accuracy and (when applicable) tune post-hoc calibration. Results are averaged across 5 seeds.

#### Fine-tuning

Unless otherwise noted, we fine-tune each backbone with identical optimization settings: Adam optimizer, learning rate 2e-5, batch size 32, 2 epochs, updating the full model. The epinet is trained for 1-2 epochs separately on the same data with the base model frozen.

#### Uncertainty methods

Temperature scaling is fit with a single temperature *T* for each 𝒟_val_ dataset (by minimizing NLL), and applied to the logits *z/T* at inference. MC-dropout enables dropout at inference (rate *p* = 0.1) and averages *K* = 10 stochastic forward passes to form 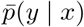. Epinet models are also evaluated with *K* = 10 epistemic samples, similarly aggregated into 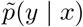.

#### Shift and OOD evaluation

We evaluate the selected model on 𝒟_test,ID_ and on shifted sets 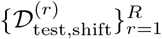, defined either by regulatory task mismatch or taxonomic novelty regimes. No tuning is performed on shifted sets. For OOD detection we treat 𝒟_test,ID_ as ID and each shifted set 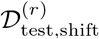 as OOD, and report AUROC using total, aleatoric, and epistemic uncertainty as the detection score separately.

### Evaluation Metrics

We assess the impact of different uncertainty quantification methods along three complementary axes: classification performance, probabilistic calibration, and out-of-distribution (OOD) detection. Predictive performance is measured directly, using classification error as a baseline for model performance.

Probabilistic calibration is evaluated using the expected calibration error (ECE, Guo et al. [20]), computed using 50 equal-mass bins:

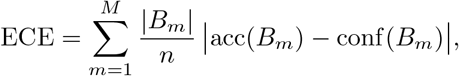

Here, predictions are partitioned into *M* = 50 equal-mass bins, and each term measures the absolute gap between empirical accuracy and average confidence within a bin. Here, lower values indicating better alignment between softmax predicted probabilities and observed frequencies. For select tasks, we additionally show reliability plots using 20 equal-width bins (shared across models to facilitate comparison), with mean curves across seeds and shaded standard-deviation bands. These plotting bins are used for visualization only and do not affect the reported ECE values. For stability, bins with fewer than 5 examples per seed are omitted from the reliability plots.

To quantify how well uncertainty signals discriminate between in-distribution and out-of-distribution examples, we treat an uncertainty score (e.g., predictive entropy or maximum class probability) as a detection statistic and compute the area under the receiver operating characteristic curve (AUROC). An AUROC of 0.5 corresponds to random detection, whereas higher values indicate that the uncertainty metric more reliably assigns higher uncertainty to OOD than to ID samples [39, 51].

## Results

### Regulatory Classification

We begin by evaluating uncertainty-aware fine-tuning on regulatory sequence classification tasks spanning promoter recognition (`promoter_all`), enhancer-type classification (`enhancers_types`), and splice acceptor prediction (`splice_sites_acceptors`). We compare four pretrained genomic backbones (see Table 1) under four inference strategies: a deterministic baseline (Ba scaled baseline (Temp scaling), Monte Carlo dropout (Dropout) epinet implementation (epinet). In addition to in-distribution (ID) task’s held-out split, we assess distribution shift via task-mismatche (Table 2). Figure 4 summarizes these trends by jointly plotting EC error for all regulatory task pairs across backbones.

**Table 1.** Genomic language models (GLMs) evaluated in this study.

| Model | Architecture | Pretraining / design (reported) |
| --- | --- | --- |
| Nucleotide Transformer v2 (500M, multi-species) [5] | Transformer | Multi-species genomic pretraining; 6-mer tokenization |
| DNABERT-2 (117M) [7] | Transformer | Multi-species pretraining; Byte-pair encoding tokenizer |
| HyenaDNA (32k small) [9] | Hyena (implicit convolution) | Long-range human genome pretraining; single character tokenization |
| CARMANIA (160k, Human) [26] | Transformer + transition matrix | Long-context human genome pretraining (GRCh38 reference); single character tokenization. |

**Table 2.**
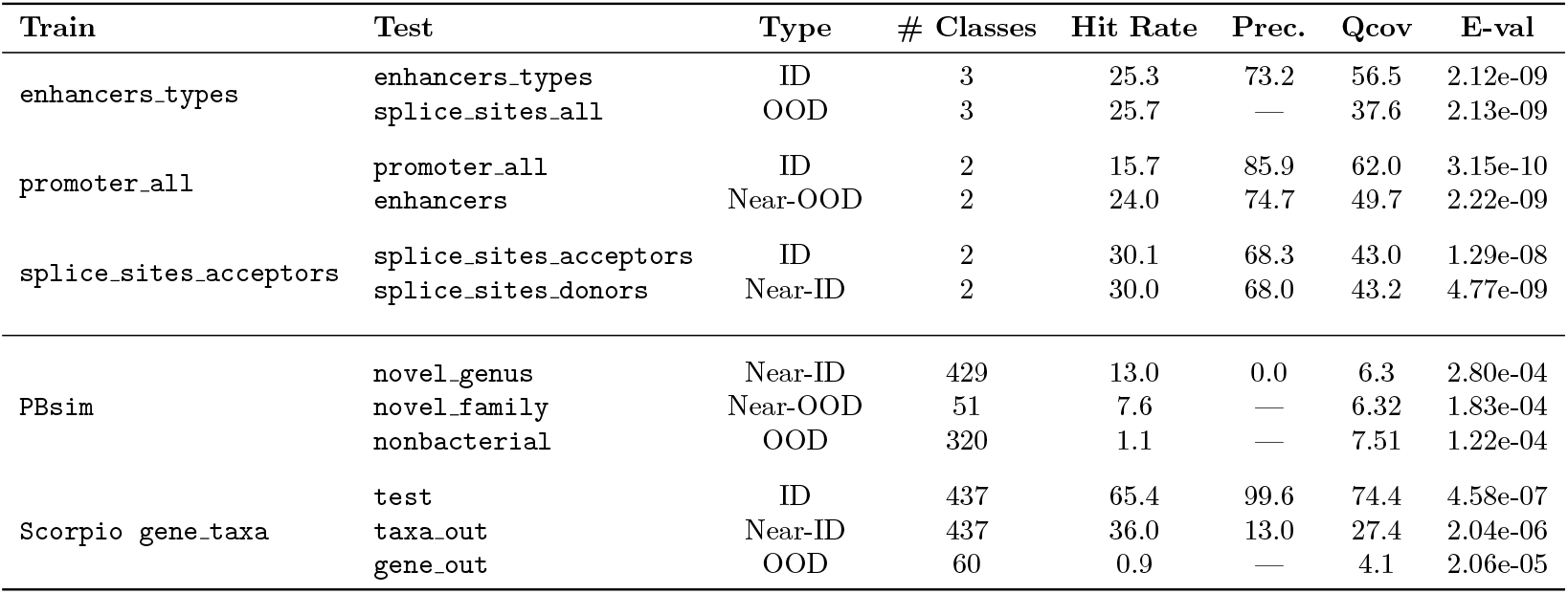
Datasets with BLAST alignment performance. (Hit rate = percentage of test sequences with BLAST alignments, qcov = query coverage, prec. = precision). Higher values indicate stronger sequence similarity between training and test sets. Lower E-val reflects that alignments are statistically significant even across regimes. Based on alignments and biological relation, datasets are assigned as **ID, Near-ID, Near-OOD**, and **OOD** categories.

**Fig 4.**
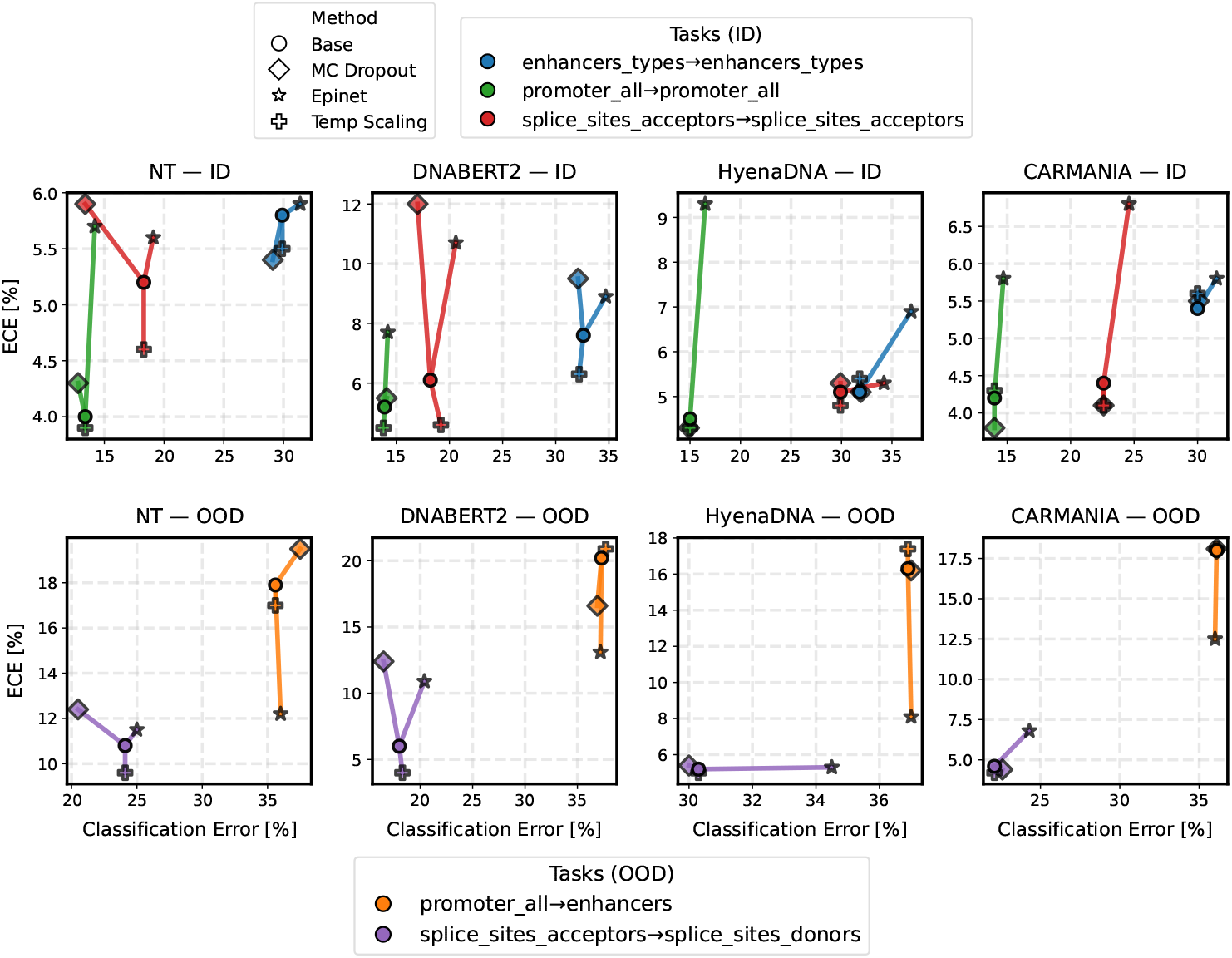
Regulatory classification performance: calibration and error. Each panel shows results for a GLM backbone (columns) and evaluation regime (rows). Points plot ECE vs. classification error, where lower is better on both axes. Colors indicate the task pair, and marker shapes indicate the uncertainty quantification method. Lines connect each uncertainty method to the corresponding Base model for the same task.

#### On ID regulatory tasks where baseline GLMs perform well, temperature scaling can further improve calibration

On ID splits, stochastic UQ variants induce only modest changes in classification error relative to Base. Across backbones and tasks, neither Dropout nor epinet provides consistent accuracy gains, and degradations are common (see HyenaDNA, column 3 Fig 4). Prediction performance on matched test distributions is largely determined by the pretrained backbone and the task difficulty, with uncertainty modeling playing a secondary role for accuracy. Any induced stochasticity from UQ generally serves to cause the models to deviate from tightly-formed decision boundaries, resulting in higher volumes of misclassifications.

Similarly, calibration on ID data is strong for all backbones, with no baseline models exceeding 8% ECE. Here, integrating Dropout and epinet yields mixed ID calibration outcomes: some settings show minor improvements, while others exhibit worse ECE than Base. In these tasks, with relatively small shifts between ID and OOD evaluations, TEMP Scalinggenerally produces the best results, achieving small improvements over the baseline in 9/12 settings. This result is expected: When training and validation datasets match the paired testing data closely, scaling the logits to that dataset will often decrease ECE and improve calibration at inference. Similar behavior can also be seen in OOD regimes that produce particularly mild shifts, namely the `splice_sites_acceptors → splice_sites_donors`, where TEMP Scaling continues to produce results that are narrowly better calibrated than the baseline.

#### Epinet consistently improves OOD calibration under regulatory domain shift

Across task-mismatched evaluations (Figure 4, OOD row), there is a clear drop-off in both classification error and calibration when compared to ID counterparts. In these conditions, epinet provides the most significant improvements in calibration among the UQ methods relative to the baseline. Under the `promoter_all` → `enhancers` shift, epinet reduces ECE for every backbone, indicating that predicted probabilities better track empirical correctness when models are evaluated away from their training distribution. This effect is clear across all models, although it is especially pronounced for HyenaDNA, where epinet halves ECE (from 16.3% to 8.1%). Figure 5 reliability plots provide a complementary view of this behavior: under ID evaluation, methods are generally similar (though the epinet can produce some underconfidence), however, under strong distribution shift, epinet more closely tracks the diagonal and reduces systematic overconfidence relative to the baseline for all models, consistent with the aggregate ECE reductions.

**Fig 5.**
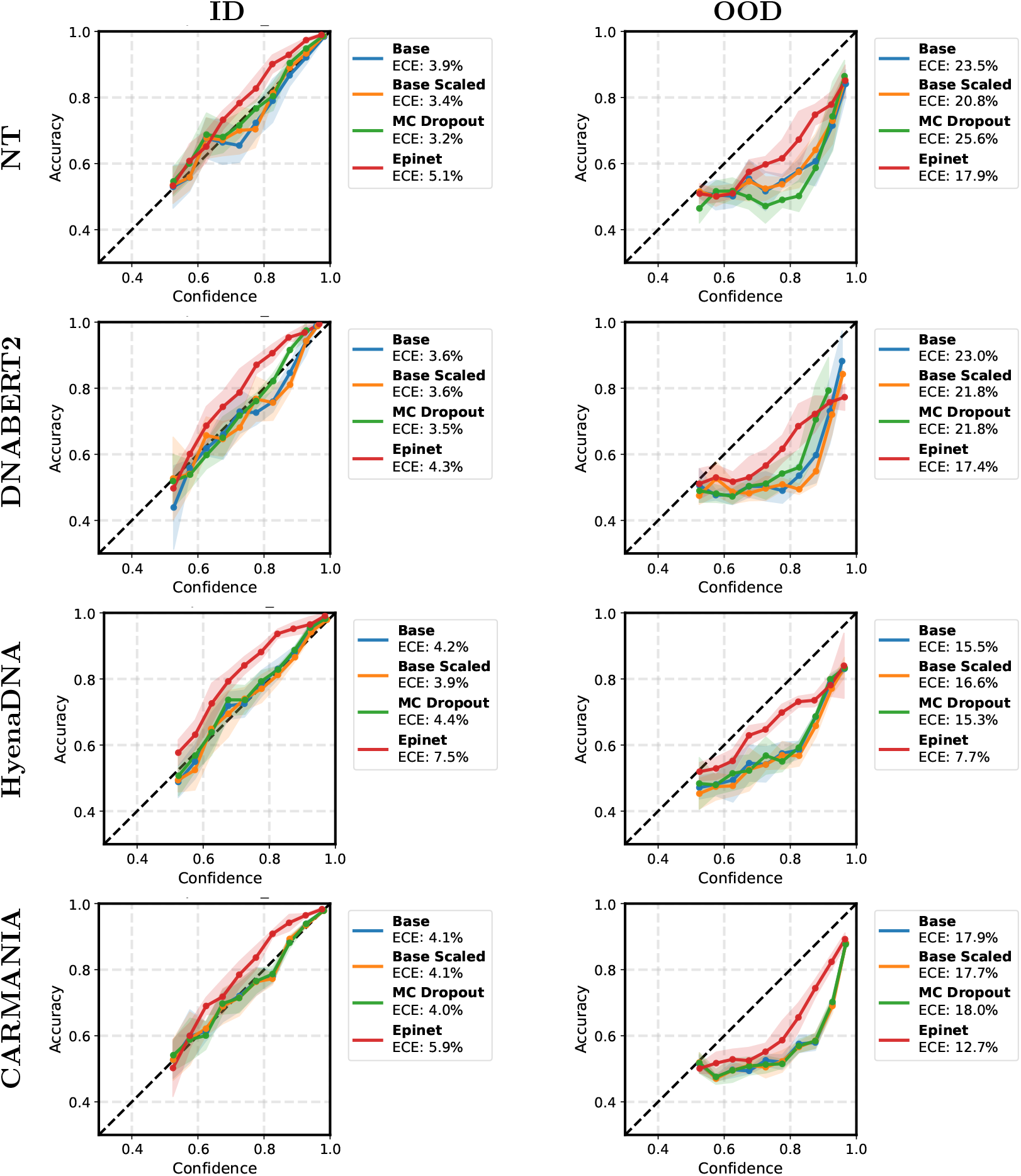
Regulatory task reliability plots. Left: ID promoter_all → promoter all. Right: OOD promoter_all → enhancers. Curves plot empirical accuracy versus predicted confidence; the diagonal indicates perfect calibration. Percentage values denote the calculated 50-bin ECE for the respective models. Under shift (right), epinet reduces systematic overconfidence relative to Base, producing a reliability curve closer to the diagonal and lower ECE.

The `splice_sites_acceptors → splice_sites_donors` evaluation produces a much more mild distribution shift. As a result, Temperature scaling is the only model to produce calibration improvements over the baseline in this setting, consistent with its performance in ID evaluations. However, while these gains are relatively steady across all GLMs, the performance gains are small overall, with a maximum ECE reduction of 2% over the base. By contrast, Dropout is overall less reliable: while it can occasionally improve calibration, it also produces degradations on the same OOD settings, and is not consistently predictable across models or datasets like Temperature scaling or the epinet. Therefore, stochastic ensembling alone does not guarantee robust uncertainty estimates.

#### Uncertainty decompositions do not reliably improve OOD detection for regulatory tasks

Assessing whether uncertainty signals can be used to discriminate OOD from ID samples across all backbones and regulatory shifts shows limited evidence that uncertainty decomposition yields a robust OOD detector in regulatory settings. Here, we compare results across singular trained models, separating the ID and OOD test sets corresponding to each trained dataset. For example - for a singular model trained on `promoter_all`, we use uncertainty scores to discriminate between samples from the `promoter_all` and `enhancers` test sets.

Relative to the softmax baseline, mean AUROC changes are small and inconsistent across models, and are often negative: for NT Transformer (Figure 6), every alternative uncertainty score reduces AUROC by a small amount on average (e.g., ΔAUROC ≈ −0.03 for Temp, ≈ −0.01 for Dropout, and larger degradations for some decomposed epinet scores), while DNABERT exhibits only a marginal average improvement for Temp (ΔAUROC ≈ +0.004) with all other variants slightly worse. HyenaDNA is near-flat, with essentially no systematic gains from decomposition, and CARMANIA is similarly mixed, showing at best small, non-generalizing changes.

**Fig 6.**
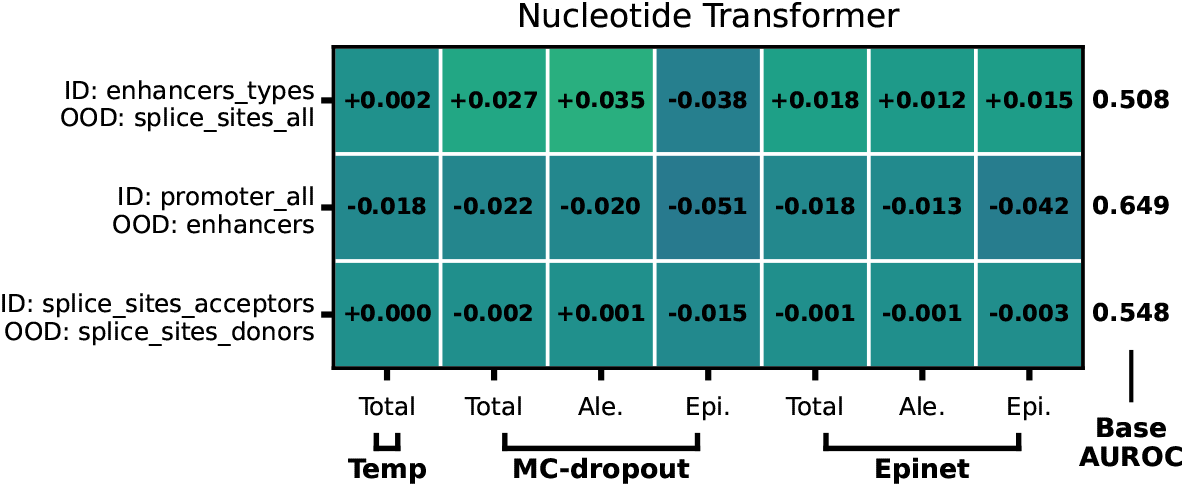
Nucleotide Transformer OOD detection (ΔAUROC vs. Base). Heatmaps report AUROC differences relative to BASE (Δ = method − `base_total`) for MC-dropout and epinet uncertainty scores (Total/Aleatoric/Epistemic). Rows correspond to ID→ OOD evaluation pairs; columns group MC-dropout and epinet.

Notably, the epistemic component AUROC is frequently lower than total uncertainty for OOD discrimination. Therefore, the epistemic score did not act like a reliable novelty detector, often ranking ID and OOD sequences less distinctly than total or aleatoric uncertainty, suggesting it was responding more to example difficulty or model idiosyncrasies than to whether a sequence came from a shifted domain. Taken together, while UQ methods can improve calibration under shift (ECE), their total and decomposed uncertainty scores do not consistently translate into improved OOD sample detection as measured by AUROC.

For the full results for OOD detection, see supplement.

### Metagenomic Classification

We next evaluate uncertainty-aware fine-tuning on metagenomic classification tasks designed to probe generalization across taxonomic structure and functional shift. We considered gene classificationon `MsAlEhR/scorpio-gene-taxa` withID(`test`), Near-ID (`taxa_out`), and OOD (`gene_out`) evaluation conditions, and simulated long reads labeled at the bacterial family level with Near-ID novelty (`novel_genus`), Near-OOD novelty (`novel_family`), and domain-level OOD shift (`nonbacterial`) (Table 2). The results of this evaluation are shown in Figure 7.

**Fig 7.**
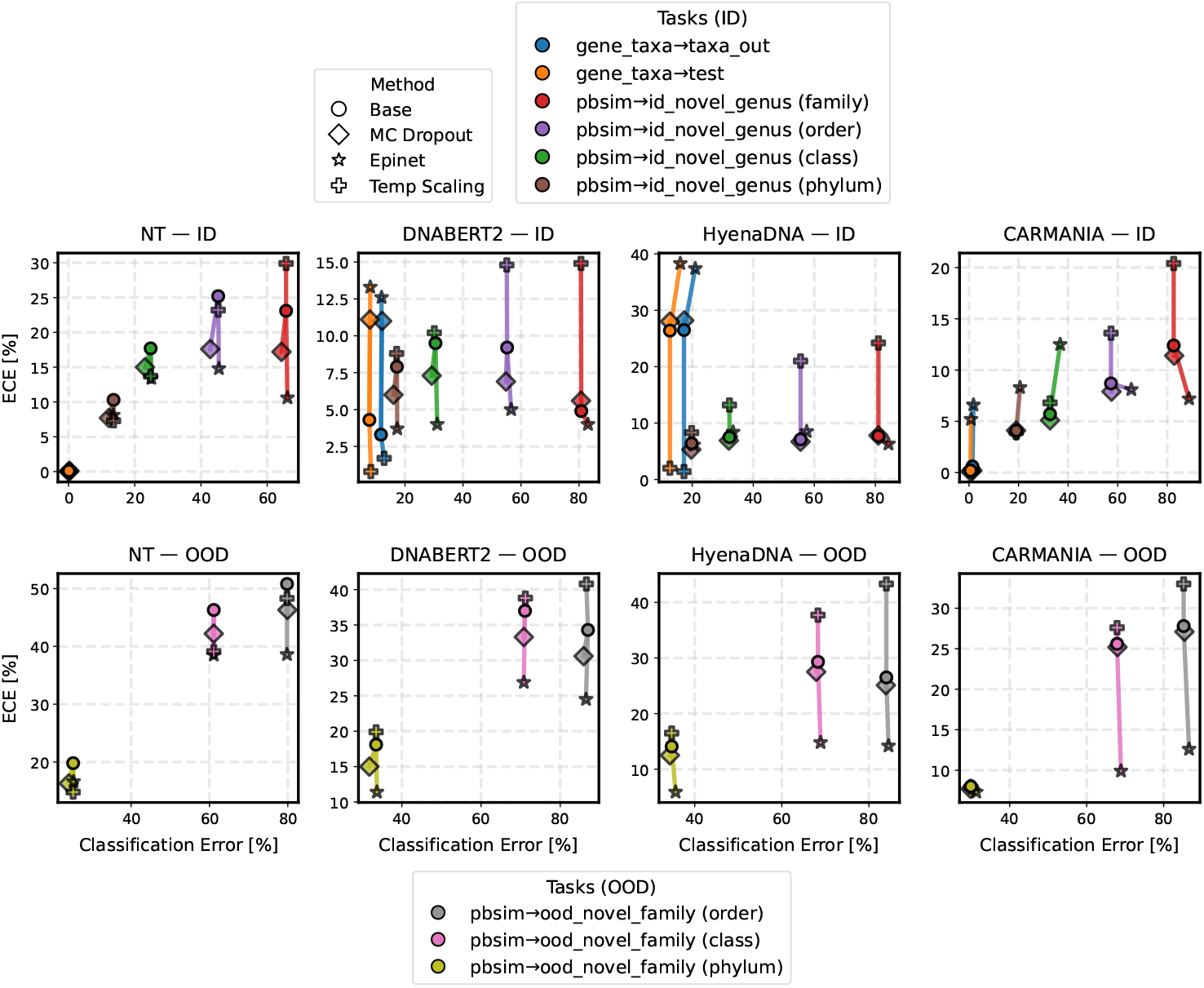
Metagenomic classification performance: calibration and error. Each panel shows results for a foundation model (columns) and evaluation regime (rows: ID vs. OOD). Points plot ECE versus classification error, where lower is better on both axes. Colors indicate the task pair, and marker shapes indicate the uncertainty method. Lines connect each uncertainty method to its corresponding Base model for the same task.

#### Simple post-hoc calibration can be extremely effective on mild ID data shifts

On the `gene_taxa` gene-class prediction task, under the majority of the ID backbone–task pairings, the baseline softmax is already well calibrated and robust to slight distribution shifts. Here, baseline models produce results with low ECE and error rates on both the entirely ID `test` evaluation as well as the slightly shifted `taxa_out` evaluation. In these regimes, additional stochasticity from approximate Bayesian models (Dropout, epinet) often yields mixed outcomes and can degrade both calibration and error, consistent with perturbing otherwise stable decision boundaries on matched distributions.

In contrast, TEMPERATURE SCALING is consistently effective on highly represented ID settings, where the held-out validation data that is used for correction is similar to the testing sets. For both datasets (`test` and `taxa_out`), temperature scaling can sharply improve calibration for models that are otherwise more poorly calibrated (e.g., HyenaDNA: 26.5% → 1.4%; BERT: 3.3% → 1.7%), and maintains already excellent calibration for NT and CARMANIA. Consistent with regulatory task findings, when an evaluation setting is well captured by the calibration data, simple post-hoc calibration is often extremely effective.

#### Temperature scaling is brittle under distribution shift

However, the same property that makes Temperature scaling attractive on ID data also makes it brittle when the test data differs from the calibration dataset. In taxonomy prediction (`pbsim`) experiments, even the ID setting (`id_novel_genus`) departs from the training data, as it shares no representative genera with the initial training. The OOD prediction introduces an even stronger shift by containing no families shared with training. In these stronger novelty regimes, temperature scaling is frequently unreliable and can substantially worsen calibration. As a result, across the majority of taxanomic prediction evaluations, Temp scaling shows elevated ECE relative to Base, raising ECE in 12/16 ID (max increase of 16.5%) and 9/12 OOD (max increase of 16.8%) tasks. These observations demonstrate a key limitation: while a single post-hoc correction can succeed under mild, well-represented shifts, they are prone to collapse when faced with significant distribution changes. This result is consistent with prior work, which has shown that temperature scaling can degrade calibration under dataset shift [34, 52].

#### Epinet provides the most consistent calibration improvements on difficult taxanomic shifts

Stronger taxonomic shifts substantially increase both error rates and miscalibration, especially as training/testing distributions become increasingly distant on lower taxanomic levels. However, this does leave room for meaningful calibration improvements to be made. Here, the epinet delivers the most consistent gains, reducing ECE relative to the baseline for every backbone, with large mean decreases for NT Transformer (−7.8%), BERT (−8.8%), HyenaDNA (− 11.6%), and CARMANIA (−10.5%). These improvements occur on challenging shifts where classification error is high (often in the ∼60–85% range), suggesting that the epinet’s primary benefit is to substantially reduce overconfidence in challenging OOD environments. Figure 8 shows reliability plots for DNABERT2, comparing ID and OOD taxonomic classification. Here, it is clear that at all taxonomic levels, and for both degrees of domain shift, the epinet is the most effective tool for decreasing overconfidence.

**Fig 8.**
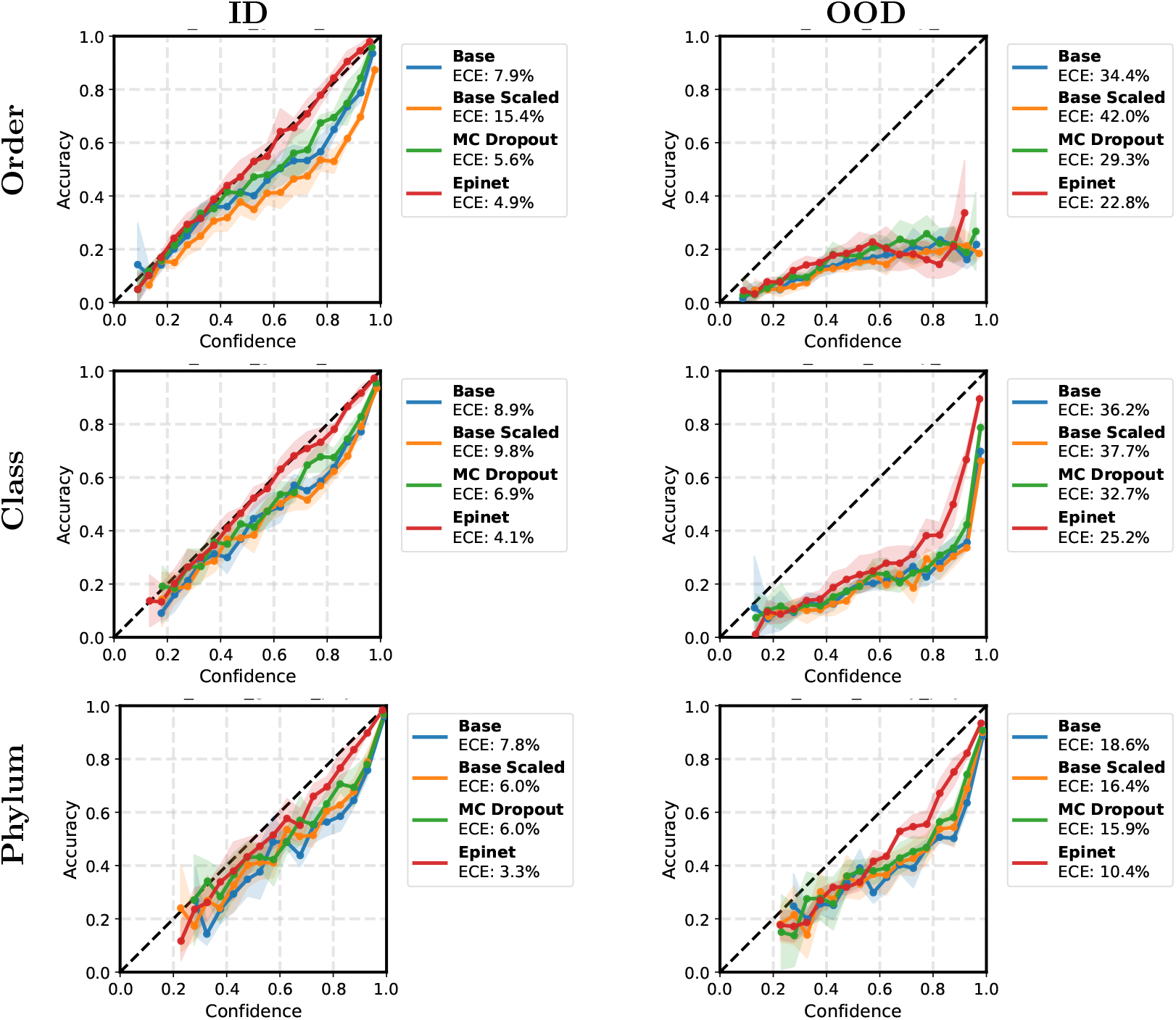
Metagenomic reliability plots at higher taxonomic ranks (DNABERT2).Left column: Near-ID evaluation (id novel genus). Right column: Near-OOD evaluation(ood novel family). Curves plot empirical accuracy versus predicted confidence; the diagonal indicates perfect calibration. Percentage values denote the calculated 50-bin ECE for the respective models Under taxonomic novelty, epinet more closely tracks the diagonal than other methods, indicating improved calibration at both ranks.

As taxonomic distribution shift becomes stronger (phylum → order classification), all models become increasingly miscalibrated on OOD data. However, epinet frequently provides the largest improvements in mitigating this overconfidence. Compared to Droupout (smaller, less uniform ECE reductions) and Temperature scaling (frequent OOD degradation), these results support epinet as the most reliable method in this study for improving calibration under significant taxonomic distribution shift.

#### Uncertainty-based OOD detection remains inconsistent

Across both metagenomic benchmarks, uncertainty scores yield mixed performance as OOD detectors, broadly mirroring the regulatory-task finding that improved calibration does not translate into strong distribution-shift detection. In many settings, AUROC differences relative to the baseline are small or negative, and there is no single method that dominates across backbones and shift types. Even when OOD detection is easy (e.g., `gene_out` for NT and CARMANIA, where AUROC is already ≈ 0.97–0.98 under Base), most UQ variants provide only marginal improvements, suggesting limited practical added value beyond the baseline signal.

A notable exception is CARMANIA on `pbsim`, where the conv epi uncertainty scores substantially improve OOD detection for challenging shifts. Figure 9 shows select tasks where the epinet provides significant gains over the baseline. In particular, for `id_novel_genus → ood_nonbacterial`, CARMANIA’s AUROC increases by 0.067 using Conv_epi total uncertainty, and by 0.161 when using the aleatoric component. Similarly, for `ood_novel_family`, AUROC increases by 0.05 (total) by 0.063 (aleatoric). While these gains do not generalize to other backbones, and decomposition into aleatoric versus epistemic components is not consistently beneficial overall, they suggest that, at least for some architectures, uncertainty estimates can capture shift-relevant structure that is not present in the baseline softmax confidence.

**Fig 9.**
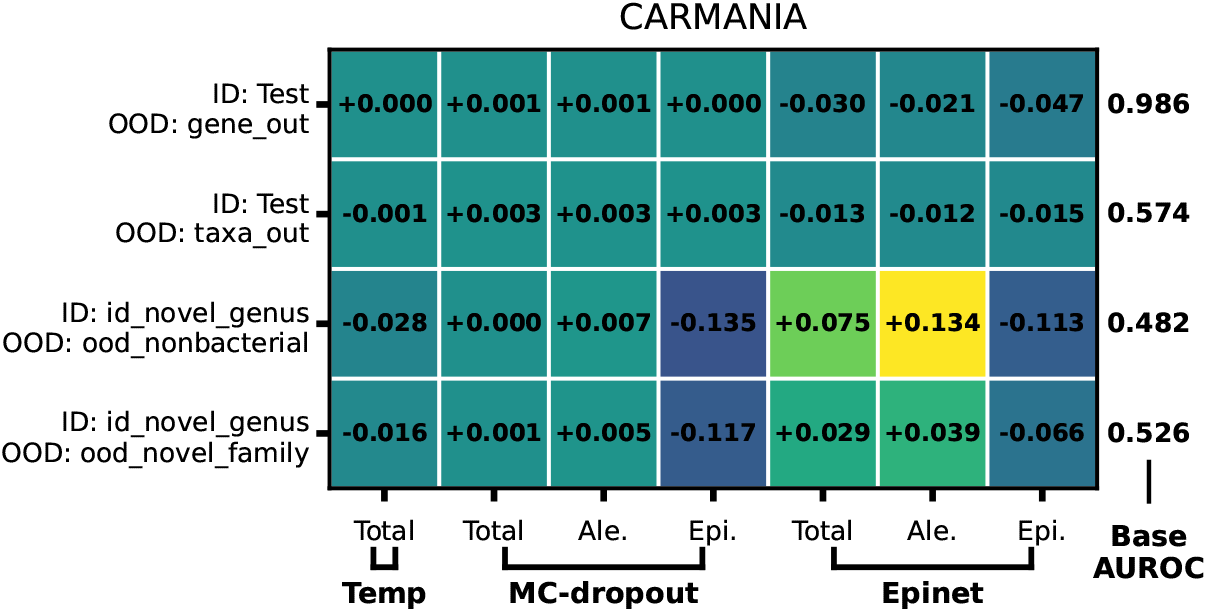
CARMANIA OOD detection (ΔAUROC vs. Base). Heatmaps report AUROC differences relative to Base (Δ = method − `base_total`) for MC-dropout and <sc>epinet</sc> uncertainty scores (Total/Aleatoric/Epistemic).Rows correspond to ID→OOD evaluation pairs; columns group MC-dropout and epinet.

For the full OOD detection results, see supplement.

### Calibration of Kraken2 and MMseqs

We evaluated calibration for two widely used bioinformatics tools (Kraken2 [53] and MMseqs2 [54]) by interpreting their native scoring outputs as confidence surrogates. For Kraken2 we compute a confidence score from k-mer match statistics (see supplement for details), because although Kraken2 provides a confidence filtering system [55], it does not output a per-prediction probabilistic confidence score, which we require for downstream comparison. For MMseqs2, we use the raw alignment-derived similarity measures percent identity (pident) and query coverage (qcov) as proxies for confidence.

Figure 10 compares these tool-based confidence surrogates to the learned DNABERT2 baseline using reliability plots under both in-distribution (ID) and out-of-distribution (OOD) conditions. Although ECE can appear favorable for Kraken2/MMseqs2 in aggregate, reliability curves show substantial departures from the identity line, particularly at intermediate score levels where calibrated probabilities should track empirical accuracy. In several settings, the MMseqs2 pident scores concentrate into a narrow high-confidence and accuracy range exclusively, leaving large regions of the confidence/probability space unoccupied. With limited coverage, ECE can be small even when the score-to-accuracy mapping is not probability-like across the range of possible values. Furthermore, even within this narrow space, the alignment of these probabilities is often significantly different from the ideal — fitting a line in the reliability plot space shows this clearly: despite low apparent ECE values, there is an inverted correlation between pident and accuracy in some cases. This observation is likely a result of higher pident corresponding to lower qcov (which despite lower ECE is visually much more reflective of confidence, particularly in promoter/enhancer identification), as shorter query coverage may result in a higher percent identity in the matched sequence. Kraken2, on the other hand generally produces better ECE and visual alignment than MMseqs2. However, this is no without cost: Kraken2 is a pure taxonomy predictor (and as such has no results for regulatory and functional predictions), and despite some alignment to the identity line, is highly variable bin-to-bin under our confidence scoring system.

**Fig 10.**
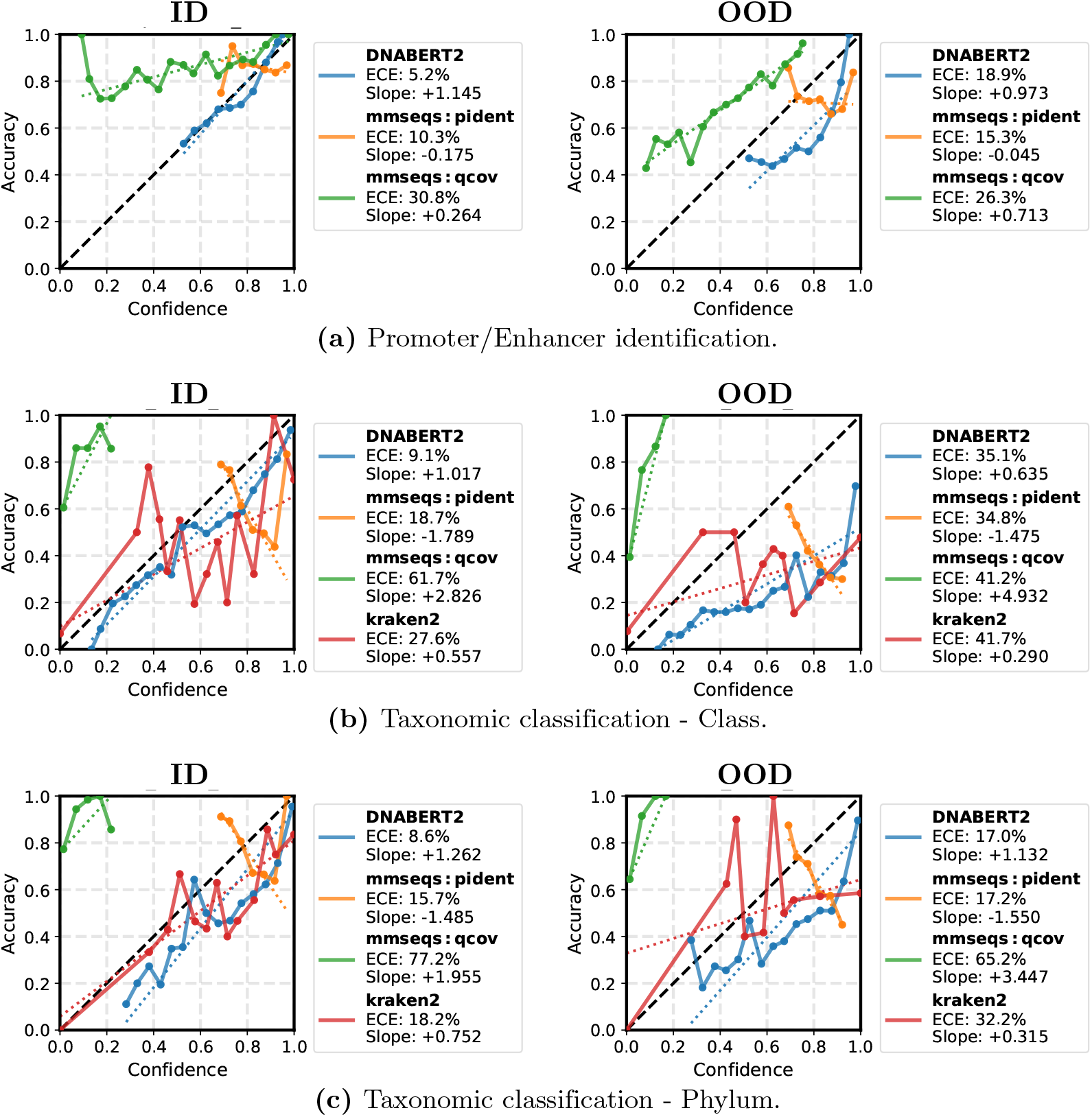
Reliability plots for tool-based confidence surrogates versus a GLM baseline. Reliability diagrams for Kraken2 confidence and MMseqs2 similarity surrogates (percent identity and query coverage), compared against the DNABERT2 softmax baseline, under in-distribution (ID) and out-of-distribution (OOD) settings for (A) promoter/enhancer identification, (B) class-level taxonomic classification, and (C) Phylum-level taxonomic classification. Points/bins show empirical accuracy as a function of reported confidence; the dashed diagonal denotes perfect calibration.

In contrast, DNABERT2 produces explicit probabilistic outputs that map more cleanly to predictive difficulty, making calibration errors more interpretable. This result demonstrates a qualitative gap between standard bioinformatics tools and modern deep learning predictors: alignment- and k-mer-based pipelines provide valuable metrics for scoring and filtering, but are not well equipped to express calibrated confidence estimates. Deep learning models, by construction, output probability distributions and can therefore be evaluated and improved using established calibration methodology [20, 34].

## Discussion

This study evaluated uncertainty quantification (UQ) for genomic language models (GLMs) across two complementary regimes: regulatory sequence classification and metagenomic prediction, each under both matched (in-distribution; ID) and shifted (out-of-distribution; OOD) evaluation. Across backbones and tasks, a consistent theme emerges: UQ methods primarily improve the quality of confidence rather than the quality of decisions. In ID settings, many model–task pairs are already reasonably well calibrated. In these regimes, stochastic approaches such as MC-dropout [19] and architectural modifications such as epinet [42] yield mixed outcomes and can degrade either calibration or error, suggesting that added stochasticity or auxiliary heads may perturb otherwise stable decision boundaries without addressing the dominant sources of predictive error.

Within the ID regime, temperature scaling is the most reliable and computationally lightweight calibration method when a representative validation set is available [16, 20, 23]. Here, it can substantially reduce systematic overconfidence while leaving classification error essentially unchanged. Notably, this benefit can extend to settings with mild shift when the calibration split captures the relevant variation (e.g., the `gene taxa` gene-class prediction task evaluated on held-out taxa with the same gene classes). However, temperature scaling is intrinsically limited by its use of a single global correction: when the relationship between confidence and correctness changes under stronger novelty, the fitted temperature may no longer be appropriate and calibration can deteriorate rather than improve [20, 23, 34].

Under distribution shift, epinet provides the most consistent improvements in calibration across both domains. In regulatory task mismatches and in metagenomic novelty regimes (including taxonomic shifts with unseen genera/families), epinet repeatedly reduces ECE relative to the softmax baseline, and it does so more uniformly across backbones than MC-dropout. Importantly, these calibration gains often occur without commensurate reductions in classification error. This separation indicates that epinet’s primary contribution is to reduce overconfidence and better align predicted probabilities with empirical correctness, rather than to substantially alter the decision boundary. For genomics applications, this distinction is practically meaningful: calibrated probabilities can support thresholding, abstention, and prioritization for downstream analysis even when raw accuracy remains unchanged [56, 57].

However, we also find that improved calibration does not imply reliable OOD detection. Across both regulatory and metagenomic tasks, AUROC-based discrimination of ID versus OOD examples using uncertainty scores (including epistemic/aleatoric decompositions [36–38]) is generally inconsistent and often close to baseline. While there are some notable exceptions–e.g. CARMANIA on metagenomic taxonomy prediction, where convolutional epinet-derived scores yield meaningful AUROC improvements on challenging shifts–Results are far from consistent, and OOD detection performance depends strongly on backbone biases and the specific form of the uncertainty estimation [18, 34, 39]. While uncertainty-aware deep learning has been widely studied for out-of-distribution detection, with successful results reported in image classification, NLP, and a range of other application domains [22, 29, 34, 39], these same results are not reflected in our genomic sequence tasks. We hypothesize that this reflects the nature of biologically realistic genomic shifts, which are often near–OOD rather than far–OOD. Despite novel function or taxonomy, unseen sequences may remain evolutionarily related or compositionally similar to the training distribution, reducing the separability of ID and OOD examples by standard uncertainty scores. This result motivates treating calibration and OOD detection as related but distinct objectives, as a method can substantially reduce miscalibration under covariate shift without producing a dependable detector of shifted inputs.

## Conclusion

We benchmarked UQ methods for four GLM backbones across regulatory and metagenomic tasks under both ID and multiple biologically motivated distribution shifts. Three conclusions follow. First, on matched ID data, calibration is often already strong and additional UQ mechanisms provide limited headroom; in this setting or similar ones, temperature scaling is an effective, low-cost calibration method when the validation distribution is representative. Second, under distribution shift, calibration degrades substantially, and epinet provides the most consistent improvements in OOD calibration across backbones and tasks, reducing overconfidence even when classification error remains high. Third, uncertainty scores and epistemic/aleatoric decompositions do not consistently yield strong OOD detection performance as measured by AUROC, although specific backbone–method combinations can occasionally improve detection in select settings. Taken together, the most robust near-term benefit of UQ for GLMs is improved probabilistic calibration, which should be implemented on a case-by-case basis depending on the degree of expected shift a model may face in practice.

## Acknowledgments

This work is supported by the National Science Foundation under Grant #2107108. We thank the University Research Computing Facility for their services, and Dr. Jim Brown for helpful discussions and for proofreading the manuscript.

